# Single-cell RNA sequencing reveals micro-evolution of the stickleback immune system

**DOI:** 10.1101/2021.12.20.473470

**Authors:** Lauren E. Fuess, Daniel I. Bolnick

## Abstract

Pathogenic infection is an important driver of many ecological processes. Furthermore, variability in immune function is an important driver of differential infection outcomes. New evidence would suggest that immune variation extends to broad cellular structure of immune systems. However, variability at such broad levels is traditionally difficult to detect in non-model systems. Here we leverage single cell transcriptomic approaches to document signatures of microevolution of immune system structure in a natural system, the three-spined stickleback (*Gasterosteus aculeatus*). We sampled nine adult fish from three populations with variability in resistance to a cestode parasite, *Schistocephalus solidus*, to create the first comprehensive immune cell atlas for *G. aculeatus*. Eight major immune cell types, corresponding to major vertebrate immune cells, were identified. We were also able to document significant variation in both abundance and expression profiles of the individual immune cell types, among the three populations of fish. This variability may contribute to observed variability in parasite susceptibility. Finally, we demonstrate that identified cell type markers can be used to reinterpret traditional transcriptomic data. Combined our study demonstrates the power of single cell sequencing to not only document evolutionary phenomena (i.e. microevolution of immune cells), but also increase the power of traditional transcriptomic datasets.

## Introduction

Pathogenic infection is a major ecological interaction that drives physiological and immune response in hosts, natural selection (4, 5), and population dynamics (6, 7). Immense natural inter- and intra-specific variation exists in organismal response to pathogens (8-10), contributing significantly disparate infection outcomes (8, 9, 12). While the consequences of variability in immunity are well documented, the underlying mechanisms which produce this variability are poorly understood. Historically, inter- and intraspecific variation in pathogenic response has been most often studied in the context of single components of the immune system (cells, genes, etc; (10, 13-16). For example, MHC II allele repertoire is significantly correlated to amphibian susceptibility to fungal pathogens; MHC heterozygosity across and within populations significantly affects pathogen resistance (17). However, recent studies have suggested that intraspecific immune variation extends beyond single components to the broad cellular structure of immune systems. Studies have documented lineage specific loss of immune cell types, as well as evolution of novel cell types in some species (18, 19). This suggests that broad scale variation in immune cell function and/or relative abundance might contribute to variation in immune responses. Still, the majority of data to this affect comes at the species level; it is unknown to what extent microevolution of immune cell identity and activity occurs within species. Understanding the extent of these processes is a necessary first step in deciphering how microevolution of immune cell types may contribute to divergence in immune response and pathogen resistance at a population level.

The immunological mechanisms underlying variable pathogen response and resistance remain particularity enigmatic in natural, non-model systems where most conclusions regarding differentiation in immunity are drawn from transcriptomic data generated from whole tissue samples (20, 21). While a powerful tool, traditional RNAseq studies condense any cell type heterogeneity within a sample to one data point. Thus, it is difficult to distinguish whether changes observed are reflective of regulatory changes in gene expression or shifting cell type abundance within the broader tissue. This is especially problematic when considering non-model species for which genetic markers of prominent cell types are lacking.

Here we leverage novel technologies in single cell RNAseq to test whether significant variation in immune cell abundance and/or function exists at the population level, potentially contributing to differentiation of immune responses. We focus our efforts on the emerging natural immunological model system, the three-spined stickleback (*Gasterosteus aculeatus*). This small fish is a tractable natural system for considering questions related to evolutionary and ecological immunology, largely due to their unique natural history. During the Pleistocene deglaciation, ancestrally anadromous populations of stickleback became trapped in newly created freshwater lakes (22). Thousands of independent lake populations have since been evolving in response to novel biotic and abiotic stimuli associated with freshwater environments for thousands of generations. This transition to freshwater exposed stickleback to many new parasites, including freshwater exclusive, cestode parasite, *Schistocephalus solidus* (23). Populations have subsequently evolved different immune traits to resist or tolerate this parasite (24). Immense variation exists between independent lake populations in susceptibility to *S. solidus* (25). Consequently, the *G. aculeatus*-*S*.*solidus* system provides a great opportunity for addressing diverse questions related to evolutionary and ecological immunity. Despite this opportunity, understanding of the broader structure of the stickleback immune system (i.e. immune cell types and functions) is limited. We conducted single cell RNA sequencing analysis to advance our understanding of immune cell repertoires and function in this important natural model system. Additionally, we leveraged the unique natural history of this species to assess questions regarding the response of immune systems to selective pressure (i.e. a novel parasite). By comparing immune cell repertoires among populations of fish which are naïve, susceptible/tolerant, or resistant to the parasite we are able to demonstrate that selection can create rapid evolutionary change in not only relative immune cell abundance, but also function (i.e. gene expression) of these immune cell types. These findings add further evidence that variation in broad immune system structure contributes to functional diversity of immunity and divergence in immune responses on a micro-scale.

## Results & Discussion

### The stickleback head kidney is comprised of eight cell types

To create a description of the immune cell repertoire of the three-spined stickleback, *G. aculeatus*, we conducted single cell RNA sequencing and associated analysis of nine laboratory-raised adult fish. Individuals were lab-raised descendants bred from wild-caught ancestors from three different populations on Vancouver Island with variable resistance to *S. solidus* (3 fish per population). These populations include one anadromous population from Sayward Estuary, which are highly susceptible to *S*.*solidus* which they rarely encounter in nature. In Gosling Lake, fish are frequently infected and tolerate rapid tapeworm growth. In nearby Roberts Lake the parasite is extremely rare, because the fish are able to mount a strong fibrosis immune response that suppresses tapeworm growth and can even lead to parasite elimination. Importantly, the fish sampled here were not infected with this cestode parasite, but instead represent constitutive population-level variability. We specifically targeted the pronephros, an important hematopoietic organ that is believed to have essential roles in the production and development of immune cells (26). Resulting libraries ranged in size from 8,119 to 19,578 cells with mean reads per cell ranging from 15,580 to 55,204 and median genes per cell ranging from 307 to 707. Following filtering (see **Methods** for details) our final data set consisting of samples ranging between 1,780 and 9,160 per library.

Analysis of resulting data revealed 24 unique clusters of cells, that could be condensed into 8 major cell types based on patterns of expression (**Figure 1; Supplementary Figure 1; Supplementary Table 1; Supplementary File 1**). These eight cell types were representative of most major vertebrate immune cell types (27): hematopoietic cells (HCs), neutrophils, antigen presenting cells (APCs), B-cells, erythrocytes (RBCs), platelets, fibroblasts, and natural killer cells (NKCs; **Supplementary Figures 2-9; Supplementary File 2**). Most of these cell types were easily identifiable based on comparison to existing data regarding vertebrate and teleost immune cell expression. For example, highly abundant neutrophils bear strong similarity to previously described teleost neutrophils, including high expression of zebrafish neutrophil marker *nephrosin* ((28)**; Supplementary Figure 3**). APCs were marked by high expression of group-specific genes involved in the presentation of antigens via the MHC II system (29, 30), and low expression of B-cell marker genes such as *cd79a* ((31)**; Supplementary Figure 5**). Also present in low abundance were a number of important minor immune cell types: platelets, fibroblasts, and NKCs; all of which were easily identifiable based on high expression of characteristic genes (**Supplementary Figures 7-9**). Interestingly NKCs were divided into two subgroups which were not easily distinguished due to low representation. One of these subgroups displayed constitutive expression of the human innate lymphoid cell (ILC) marker gene, *rorc* (32), as well as high expression of *runx3*, which modulates development of ILCs (33), providing some support that this subgroup was comprised of putative fish ILCs. Conspicuously absent were putative T-cells. This can likely be explained due to the nature of the pronephros, which is believed to operate similarly to mammalian bone marrow (34-36). Consequently, T-cells are likely only transiently found in this organ, perhaps only early in life.

**Figure 1:**
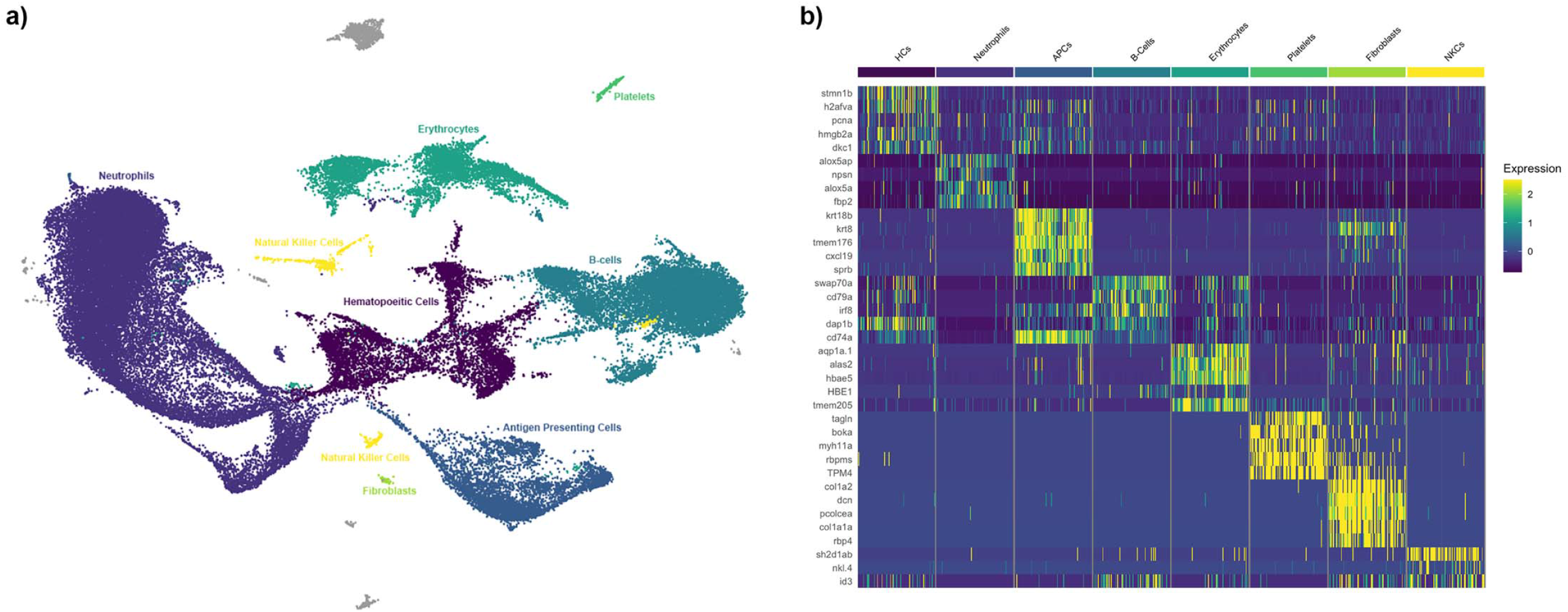
**a)** UMAP projection of head kidney cells generated from combining all 9 samples. Each point represents a single cell. Cells are color-coded by their cluster and annotated cell type. Cells are shown grouped into major cell type clusters based on distinguishing genes. For original cluster assignments see **Supplementary Figure 1 & Supplemenatry Table 1. b)** heatmap of the top five annotated distinguishing genes per cluster. Scaled expression, generated using the Seurat R package is displayed for each gene. Cells are grouped by type, genes are listed in order of significance. Only 3 annotated genes were significant for the NKC cluster.

### Stickleback erythrocytes express a variety of immune genes

In teleosts, unlike mammals, red blood cells are nucleated and genetically active (37). A large, heterogenous group of cells with high expression of hemoglobin-associated genes were identified as putative erythrocytes. Interestingly these cells also had high expression of a number of immune genes characteristic of both neutrophils and B-cells (**Figure 2**). Previous findings have indicated that teleost RBCs have diverse roles in the regulation of host immunity (38, 39). For example, it is well documented that teleost RBCs contribute to antiviral immunity (38, 40, 41). Preliminary evidence suggests they also can phagocytose and kill bacterial pathogens (42) and even yeast (43). However, our results suggest further refinement of these functions. Clustering analysis shows two distinct subgroups of RBCs, dividing based on similarity to either myeloid (neutrophil) or lymphoid (B-cells) type cells (**Figure 2**). Thus, while previous studies have both characterized myeloid type functions (38, 41, 42) and document interactions with lymphoid cells (44), this is the first evidence for diversification of RBCs into distinct subgroups, each serving a particular immunological role. Further study is needed to improve understanding of the distinct roles of these two subtypes and their broad roles in fish immunity.

**Figure 2:**
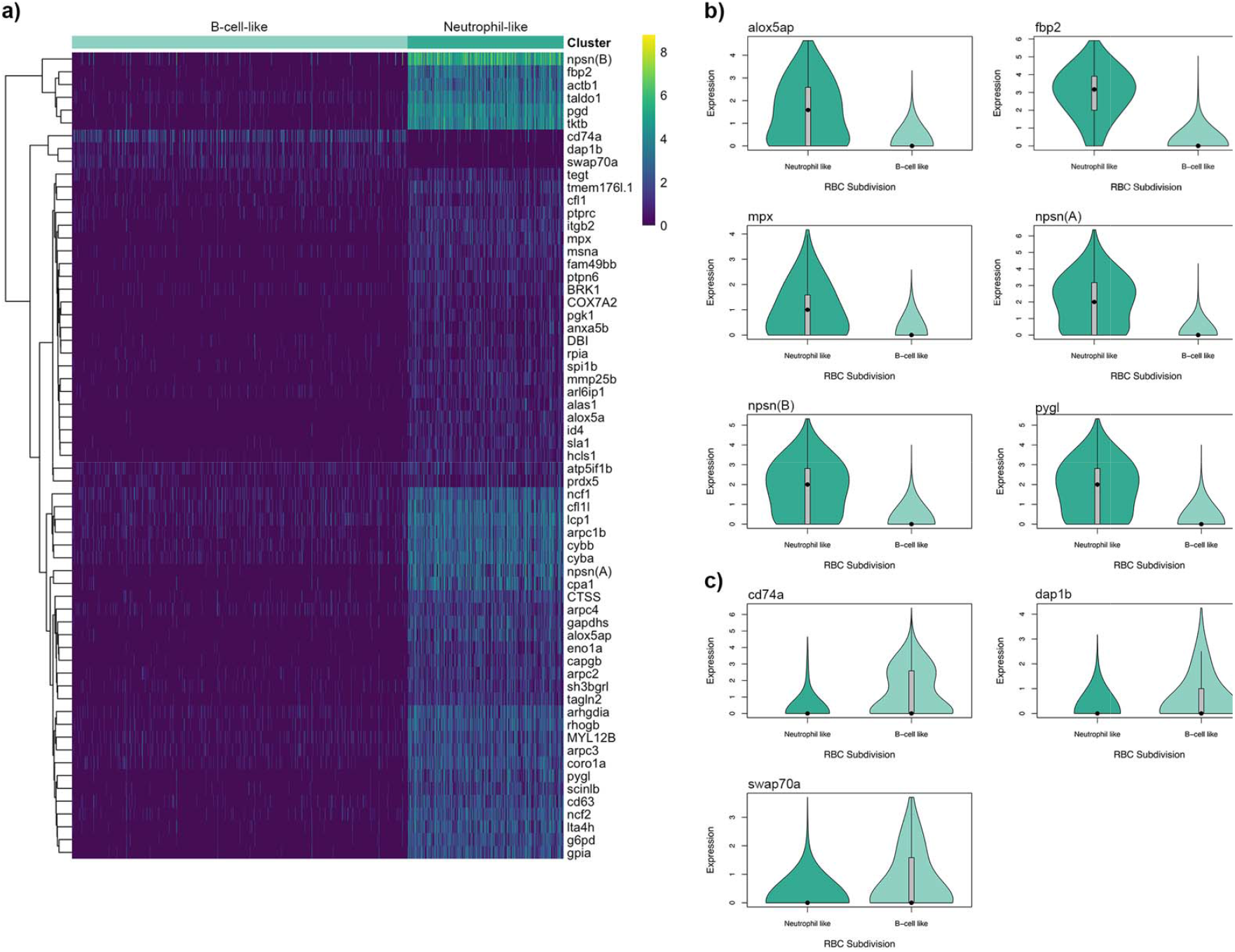
Differential expression of immune genes among the two identified RBC subgroups (neutrophil like and B-cell like). **A)** heatmap of log normailzedexpression of annotated B-cell and Neutrophil marker genes which were significantly differentially expressed between the two RBC subgroups (mitochondrial and ribosomal genes excluded). Heatmap generated using the pheatmap package in R. **B)** Violin plot of log normalized expression of significantly differentially expressed neutrophil marker genes among the two subgroups of cells. **C)** Violin plot of log normalized expression of significantly differentially expressed B-cell marker genes among the two subgroups of cells.

### Two groups of B-cells are identifiable: resting and plasma B-cells

A large group of cells uniquely expressing *cd79a, swap70a*, and a number of putative immunoglobulin genes, was identified as putative B-cells (**Figure 1**). This group was comprised of three sub-clusters (original clusters 11, 12, 13; **Supplementary Figure 1**), two of which (cluster 12 and cluster 13) were readily distinguished by expression patterns (**Supplementary Figure 10**). The smaller of the two sub-clusters (cluster 13) had considerably higher expression of immunoglobulin genes as well as *X-box binding protein 1* (*xbp1*) and associated proteins, key markers of plasma cells in mammals (45). Thus, we concluded that these two groups likely comprised of resting B cells (cluster 12) and activated/plasma B cells (cluster 13). Previous work has documented the diversification of fish B cells into antibody secreting cells upon immune stimulation (46). Furthermore, studies have indicated that antibody-secreting cells (including plasma cells and plasmablasts) constitute a stable subpopulation of cells in the head kidney of other fish species. Interestingly though, low levels of resting B-cells in the head kidney have been documented in salmonids, which is contrary to our preliminary findings here (47). High levels of resting B-cells are characteristic of tissues involved in inducible responses to immune challenge; typically the blood and spleen in teleost fish (47). However, it is possible that some fish lineages may have evolved more plasticity in head kidney function as part of an inducible immune response. Further characterization of B-cell subpopulation in other tissue types from *G. aculeatus* will provide insight regarding the lineage-specific roles of various lymphoid tissues in immunity.

### Isolated populations of stickleback vary significantly in cell type abundance

The nine fish sampled for our scRNAseq analysis were representative of three isolated and genetically divergent populations. These three populations, Roberts Lake, Gosling Lake, and Sayward (anadromous) have been well documented to vary considerably in their immune responses to a common freshwater parasite, *Schistocephalus solidus* (25). The marine population is evolutionarily naïve to the parasite, which does not survive brackish water, and consequently is readily infected and permits rapid cestode growth. Both Gosling and Roberts Lakes are more resistant to laboratory infection than their marine ancestors, but the more resistant Roberts lake population significantly suppresses cestode growth and is more likely to encapsulate and kill the cestode in a fibrotic granuloma (25, 48). Consequently, we divided our samples based on population and compared both immune cell relative abundance and within-cell-type expression across these three populations. Comparing the three populations, we find significant variation in abundance in every cell type except fibroblasts (**Supplementary Table 2**; **Figure 3**). Roberts Lake fish, which are most resistant to the parasite, had considerably more neutrophils and platelets, but significantly less NKCs, RBCs, and B-cells than the other two populations. Sayward fish, which are anadromous and naïve to the parasite, had the highest abundance of APCs, B-cells, and RBCs.

**Figure 3:**
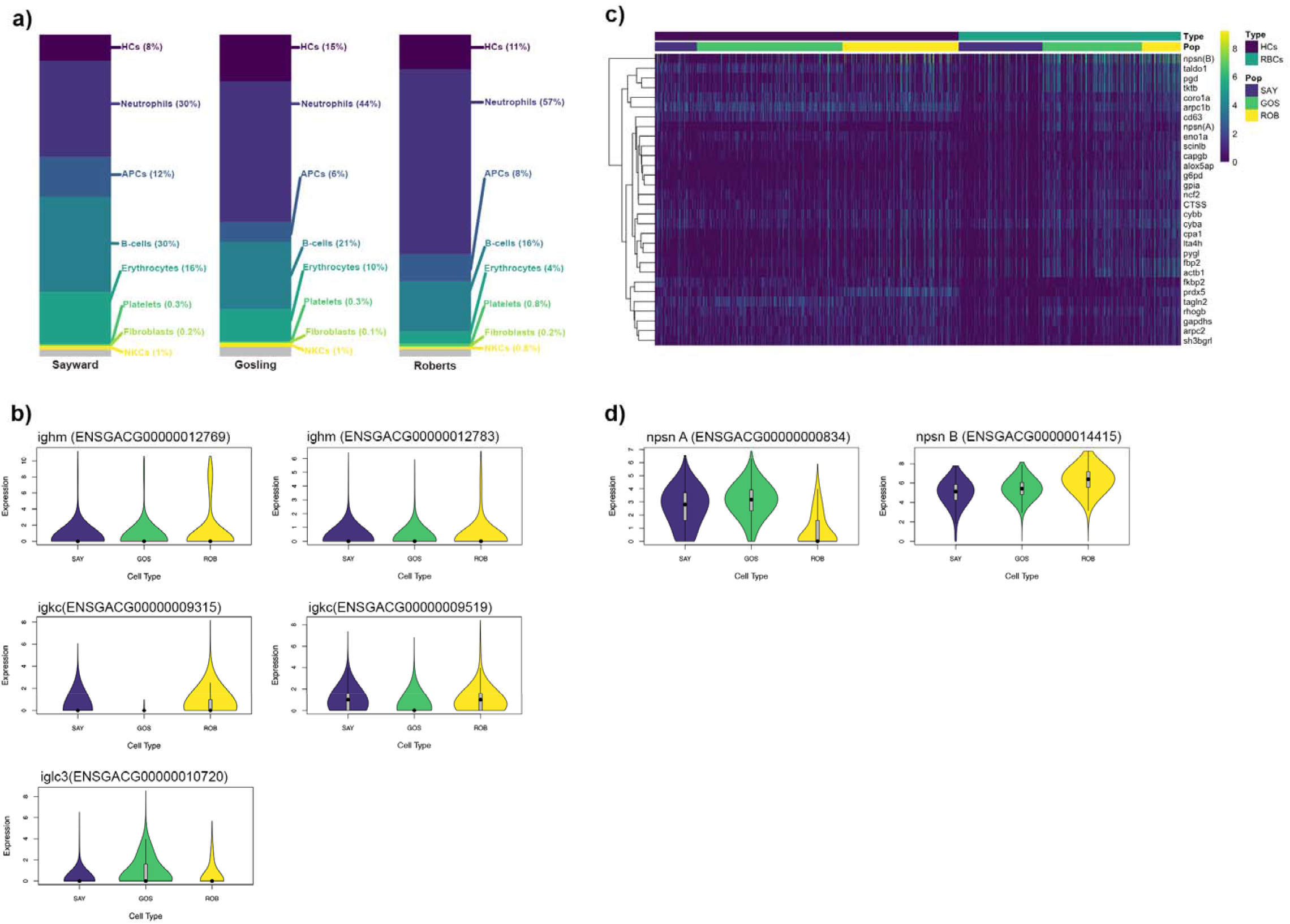
Summary of variation in immune cell subtype abundance and expression across three sampled populations of *G. aculeatus*. **a)** bar graph representing relative abundance of each of the eight major cell types within each of the three sampled populations. **b)** comparison across populations of expression of B-cell specific expression of putative immunoglobulin genes. All three genes were significantly differentially expressed between two or more populations. ^*^ indicates significantly different values. **c)** heatmap of expression of neutrophil marker genes in both the hematopoietic cells and erythrocytes. Columns are clustered by cell type and then population. Genes are clustered by similarity of expression profile using base algorithms from the pheatmap package in R, **d)** comparison of neutrophil-specific expression of the two copies of nephrosin across the three sampled populations ^*^ indicates significantly different values

Much of this observed variation in immune cell type abundance may be related to natural variation in parasite resistance. For example, Roberts lake fish had higher abundance of both neutrophils and platelets, which may contribute to enhanced resistance to helminth parasites. Neutrophils and other granulocyte cells such as eosinophils are important components of the initial innate immune response to helminths and other parasites (49, 50). Platelets, specifically thrombocyte-derived compounds, are important mediators of fibrotic responses (51, 52), and fibrosis is a major part of Roberts Lake sticklebacks’ response to *S. solidus* infection (48). Consequently, enhanced abundance of both neutrophils and platelets in ROB fish likely allows for quick induction of resistance phenotypes (i.e. fibrosis; (53) and other immune responses which result in the efficient elimination of the parasite. It should be noted that the lack of variation in fibroblast abundance among populations is not unexpected; while platelets are normally originate in hematopoietic tissues, like the head kidneys (54), fibroblasts are usually stimulated at sites of damage (55), which is in the peritoneal (body) cavity for the *S. solidus* parasite.

Combined, the differences in relative abundance of immune cell types observed among our three populations of fish are indicative of micro-evolution in response to parasites. Because the fish used in this study were lab-raised in a shared environment, these between-population differences likely reflect heritable differences that evolved since the populations were founded. Roberts Lake fish, which evolved resistance to the helminth parasite *S. solidus*, is characterized by marked increases in immune cell subtypes which (in mice) are known to contribute to helminth responses. Thus, our results suggest that evolution of resistance to a parasite may not only occur on the gene level, but that resistance may also be the result of selection for broad-scale shifts in baseline immune cell type abundance.

### Expression of each cell type varies among populations

In contrast to the significant variation in relative abundance of immune cell types between the three sampled populations, we found modest signatures of among-population variation in expression profiles within cell types (**Supplementary File 3**). Most notable was variation in expression of immunoglobulin-like genes in B-cells (**Figure 3**). Despite having significantly fewer B-cells in Roberts Lake fish, their B-cells exhibited higher average expression of immunoglobulin-type genes per cell. This may be a compensatory method as B cell production of immunoglobulin is an essential component of response to helminth infection (47). Indeed, higher expression of immunoglobulin genes by ROB B cells is likely the result of a significantly higher relative abundance of putative plasma B cells in ROB fish (compared to resting B-cells). ROB fish had higher proportions of plasma cells generally, and as a subset of B cell population than both GOS and SAY fish (Chi-squared test; *padj* < 0.001). Helminth-protective T_H_2-type immune responses induce expansion of plasma cells producing IgE (56). Thus, a higher constitutive abundance of plasma-type B-cells in ROB fish may contribute to enhanced resistance to *S. solidus* parasites. Again, here our results indicate that micro-evolution of immune cell subtype abundance may significantly contribute to evolution of parasite resistance.

Finally, patterns of expression of neutrophil-associated markers also varied significantly across populations. Both HCs and RBCs in Roberts had significantly higher expression of neutrophil marker genes (**Figure 3**). This is likely the result of enhanced overall investment in neutrophil-like cells in Roberts fish, which may support a quick initial response to invading parasites (49, 50). Perhaps most interestingly, we observed population-specific, preferential expression of what is presumably duplicated copies of the important zebrafish neutrophil marker gene, *nephrosin* (*npsn*). We identified two highly similar genes annotated as *npsn*, both of which were significant markers of neutrophils, however one gene was preferentially expressed by Roberts fish, while the other was expressed higher in Gosling and Sayward neutrophils (**Figure 3**). Sequence comparison of these two gene copies revealed that while highly similar to zebrafish *npsn*, there are several species-specific, and copy-specific amino acid substitutions in the sequences, suggesting potential neofunctionalization (**Supplementary Figure 11**). Neofunctionalization of one copy of this gene may be the result of co-evolutionary selective pressure. While neofunctionalization of parasite virulence genes has been recorded in the past (57, 58), this is to our knowledge the first evidence of neofunctionalization potentially contributing to host resistance.

### Insights from scRNAseq analyses improve interpretation of past traditional RNAseq studies

The scRNAseq data allowed us to confidently identify a suite of genes which are markers of each of these putative 8 cell types (**Supplementary File 2**). Using these new candidate marker genes, we can re-evaluate findings of past RNAseq studies to understand the relative contributions of changes in gene expression versus changes in cell abundance. Specifically we leveraged these markers to re-interpret results from two previous studies for which we had both traditional RNAseq expression data and flow cytometry data coarsely estimating granulocyte to lymphocyte relative abundance using forward and side-scatter gating (2, 59). The first, and larger, of the two studies investigated variation in constitutive and induced immune response to parasite infections in laboratory reared F2 fish (59). Within this data set, granulocyte and lymphocyte frequencies are, respectively, correlated to expression of both putative granulocyte markers (*nephrosin* B, transcript 1; pearson correlation, *p* < 0.001, *r* = 0.3904) and lymphocyte markers (*cd79a*; pearson correlation, *p* < 0.001, *r*= 0.4569). The second, smaller, study conducted a similar experimental parasite infection of laboratory reared F1 fish (2). Within this study, these correlations are less significant for lymphocytes (Pearson correlation, p = 0.016, *r* =0.25), and both non-significant and trending in the opposite direction for granulocytes (pearson correlation, p = -0.17, *r* =0.12; **Figure 4**). These inconsistencies are likely due to the nature of our correlative data. Flow cytometry grouped cells into two large bins: granulocytes and lymphocytes. Thus, finding two markers that accurately correlate to these broad groups across experiments is difficult, particularly for diverse granulocytes. Still, these findings suggest that variation in expression of cell markers identified here may be reflective of changes in abundance of immune cell types. We believe that further validation will demonstrate that this data provides a powerful new resource that will increase the interpretive power of traditional RNAseq analyses.

**Figure 4:**
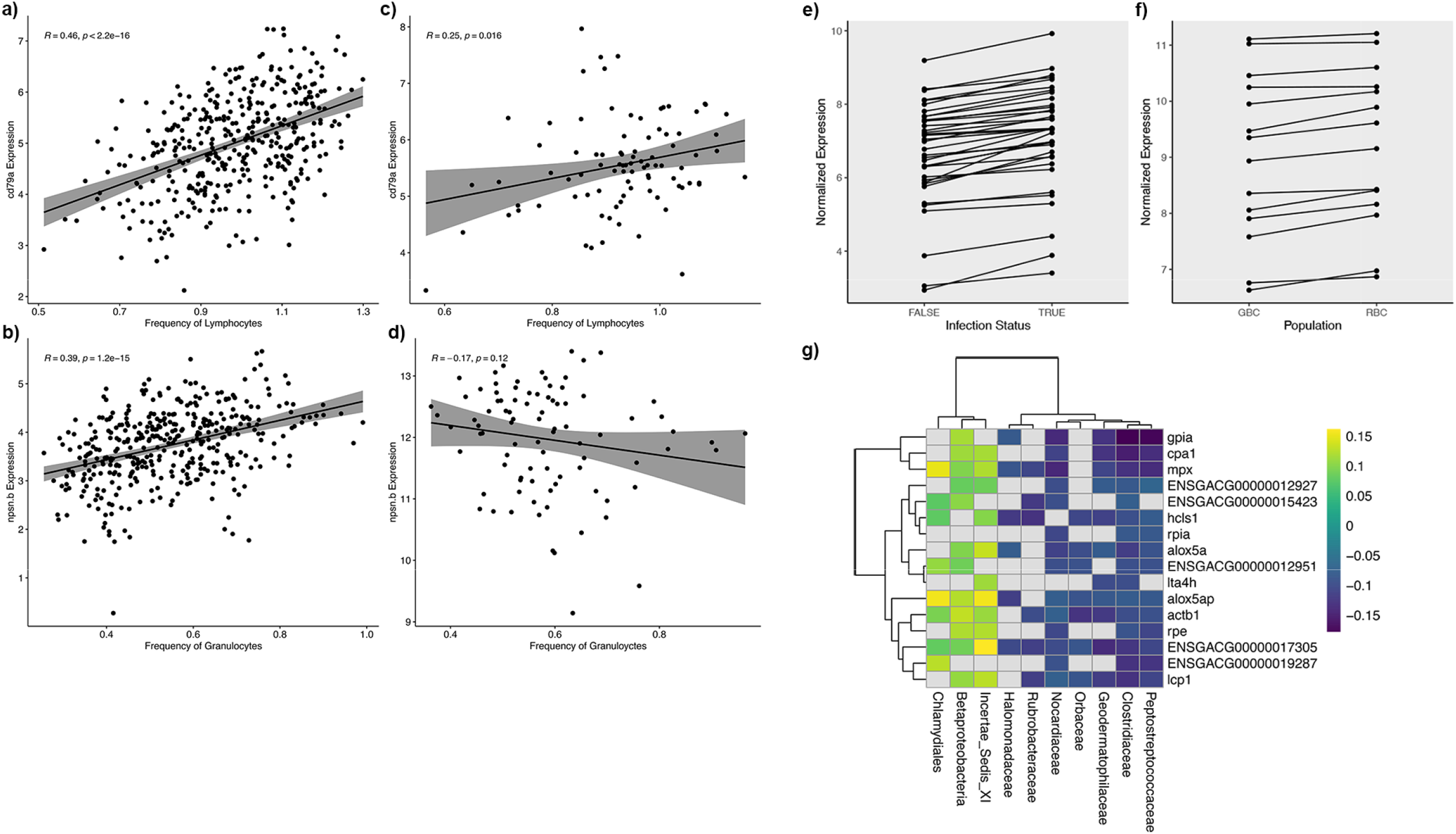
Evaluation of applicability of identified markers to past traditional RNAseq datasets **a-d)** pearson correlations between expression of identified lymphocyte (cd79a) or granulocyte markers (npsn.b) and normalized lymphocyte or granulocyte frequency (detected by flow cytometry) in our two previous transcriptomic studies sets (**a-b**; (1), (**c-d**; (2, 3). For all correlation plots regression line is shown in black and shading indicates 95% confidence intervals. **e)** patterns of differences in gene expression of identified APC in uninfected vs. infected fish **(f)** patterns of differences in gene expression of identified B-cell markers in parasite susceptible (GBC) vs. parasite resistance (RBC) fish; all data shown in **e-fx** corresponds to genes which were significantly differentially expressed in a previous traditional RNAseq study **(REF). g)** heatmap of significant correlations (tau) between gene expression of identified neutrophil markers and abundance of specific microbial taxa. Non-significant correlations are displayed in grey. Data taken from a previous correlative analysis of traditional RNAseq data (11).

Assuming that changes in expression of these markers is at least in part due to changes in their respective cell type, we can now glean more insight regarding the cellular changes in response to infection of *G. aculeatus* by *S. solidus* by re-examining previous datasets. Consequently, we applied the markers generated here to reinterpret results from the two studies of response experimental parasite infection in laboratory reared F1 and F2 fish (2, 59). In each case we conducted Chi-squared tests to detect over-representation of cell markers (generally or specific cell type) among significantly differentially expressed genes. In the case of groups where significant over-representation was detected, we conducted a proportion test to detect statically significant skew in the directionality of differential expression. In the smaller study of response of laboratory reared F1 fish, we observed few significant patterns of biological interest (2); **Supplementary Table 3**). However, in our larger dataset (F2 fish) we noticed significant over-representation of APCs and B-cell marker genes among the genes differentially expressed as a result of infection or between populations respectively (59); **Figure 4, Supplementary Table 3**). Markers of APCs were not only significantly over-represented, but also exclusively increased in response to infection. Alternatively activated macrophages are known to play key roles in response to helminth infection, including mediating inflammatory responses (60, 61). B-cell markers were expressed a higher levels in susceptible back-crossed fish compared to resistant back-crosses, consistent with analysis of scRNA data presented here.

Finally, we also considered results from correlative analyses of associations between gene expression in F2 fish, and gut microbiome composition (11). Here we observed significant over-representation of markers of neutrophil, B-cell, and fibroblast cells among lists of genes significantly correlated to abundance of specific microbial taxa in the gut (**Supplementary Table 3**). Neutrophils demonstrated the most consistent patterns of association with microbial taxa abundance, with some microbial taxa demonstrating strongly significant positive or negative associations with many neutrophil markers (**Figure 4)**. Neutrophils and gut microbiota are believed to be functionally linked, with gut microbiota regulating components of neutrophil activity and vice versa (62). Our findings suggest that specific microbiota have systemic effects on the proliferation of (or lack thereof) neutrophils in hematopoietic organs. In sum, the markers discovered here provide new power to interpret traditional RNAseq data and begin to disentangle relative contributes of changes in gene expression versus changes in cell type abundance. These results point to the value of small-sample scRNAseq in guiding reinterpretation of new or existing large-sample bulk-tissue transcriptomic data.

## Conclusions

Here we present a robust analysis of the contributions of variation in immune system structure (relative cell type abundance and function) to observed variation in immune response between populations of fish. While numerous previous studies have suggested that shifts at the genetic level contribute to variation in immune response (10, 12, 48, 63), our study is the first to look at this variation among natural populations at the cellular scale. Using single-cell RNAseq analyses, we demonstrate that independent populations vary significantly in both abundance and expression patterns of immune cell types. Furthermore, our data suggest that this variation may be the result of micro-evolution of immune cell repertoires in response to biotic stimuli (i.e. a novel parasite). This is, to our knowledge, the first evidence that rapid evolution of immune cell repertoires among populations both occurs, and potentially contributes to variation in immune response and infection outcome. Our results add to the growing body of evidence that suggests that the immune system may be much more malleable than once thought. Furthermore, these findings provide compelling rationale for further studies investigating adaptability of immune system structure within and between species in response to eco-evo feedbacks. Also notably, our findings present the first description of prominent immune cell types in an important ecological and evolutionary model species. This provides new cell marker resources that can be used to streamline further immunological studies and provide new insight into traditional RNAseq studies. In sum, or work not only adds strong evidence suggesting that micro-evolution of immune cell repertoires contributes to variation in immune response, but also provides a robust new tool for researchers utilizing the stickleback system as a model of evolutionary and ecological immunology.

## Methods

### Sample Collection & Processing

Single cell libraries were generated from head kidneys of laboratory reared F1 stickleback from three populations on Vancouver Island in Brittish Columbia (Sayward Estuary, Roberts Lake, Gosling Lake). Reproductively mature fish were collected at each location using minnow traps. Gravid females were stripped of their eggs, which were then fertilized using sperm obtained from macerated testes of males from the same lake. Fish were collected with permission from the Ministry of Forests, Lands, and Natural Resource Operations of British Columbia (Scientific Fish Collection permit NA12-77018 and NA12-84188). The resulting eggs were shipped back to Austin, Texas, hatched, and reared to maturity in controlled laboratory conditions. At approximately 2-3 years of age, fish were transferred to aquarium facilities at the University of Connecticut. At the time of sampling, fish ranged from 3 (Sayward and Gosling) to 4 (Roberts) years of age.

We generated single cell suspensions from the pronephros (head kidney) three fish from each population (Sayward, Roberts, Gosling). Fish were humanely euthanized one at a time, and their head kidneys immediately extracted. Dissected head kidneys were placed in 2mL of R-90 media (90% RPMI 1640 with L-Glutamine, without Phenol red; Gibco) in a sterile 24-well plate on ice. Tissue was then physical dissociated using a sterile pipette tip. The resulting slurry was then strained through a 40μm nylon filter. An additional 2mL R-90 was added to the resulting suspension. Cells were then spun at 440*g* for 10 minutes at 4°C. The supernatant was removed, and cells were resuspended in 2mL R-90. Cells were spun once more time, and the resulting supernatant replaced with 1 mL R-90. Cell suspensions were then transported on ice to the Jackson Lab facility in Hartford, Connecticut where samples were prepared for sequencing and sequenced within 6 hours of initial sample collection.

### Single Cell Library Preparation and Sequencing

Cells were washed and suspended in PBS containing 0.04% BSA and immediately processed as follows. Cell viability was assessed on a Countess II automated cell counter (ThermoFisher), and an estimated 12,000 cells were loaded onto one lane of a 10x Genomics Chromium Controller. Single cell capture, barcoding, and single-indexed library preparation were performed using the 10x Genomics 3’ Gene Expression platform version 3 chemistry and according to the manufacturer’s protocol (#CG00052, (64). cDNA and libraries were checked for quality on Agilent 4200 Tapestation, quantified by KAPA qPCR, and sequenced on an Illumina sequencer targeted 6,000 barcoded cells with an average sequencing depth of 50,000 read pairs per cell. Three initial libraries (1 per population) were sequenced on individual lanes of a HiSeq 4000 flow cell; all other libraries were sequenced on a NovaSeq 6000 S2 flow cell, each pooled at 16.67% of the flow cell lane.

Illumina base call files for all libraries were converted to FASTQs using bcl2fastq v2.20.0.422 (Illumina) and FASTQ files were aligned to reference genome constructed from the v5 G. aculeatus assembly and annotation files available at https://stickleback.genetics.uga.edu/ (65). Briefly, annotations from Ensembl (release 95) were combined with repeat, Y chromosome, and revised annotations from Nath et al. using AGAT (0.4.0) (66), and a STAR-compatible reference genome was generated Cell Ranger (v3.1.0, 10x Genomics) using these annotations and the v5 assembly from Nath et al. The Cell Ranger count (v3.1.0) pipeline was used to construct cell-by-gene counts matrix for each library, subsequently analyzed using Scanpy 1.3.7 (67) and the Loupe Cell Browser (10x Genomics).

Each counts matrix was individually subjected to quality control filtering, such that cells with more than 35,000 UMIs, fewer than 400 genes, more than 30% mtRNA content, and more than 1,000 hemoglobin transcripts were discarded from downstream analysis. The nine filtered counts matrices were concatenated, normalized by per-cell library size, and log transformed. The expression profiles of each cell at the 4,000 most highly variable genes (as measured by dispersion (64, 68) were used for principal component (PC) analysis and subsequently batch corrected using Harmony (69). The batch corrected PCs were utilized for neighborhood graph generation (using 25 nearest-neighbors) and dimensionality reduction with UMAP (70). Clustering was performed on this neighborhood graph using the Leiden community detection algorithm (71). Subclustering was performed on a per-cluster *ad hoc* basis to separate visually distinct subpopulations of cells. This UMAP embedding and clustering metadata were then imported into the Loupe Cell Browser (generated using Cell Ranger aggr (v3.1.0)) for interactive analysis.

### Cluster Identification

Once data (UMAP embedding and clustering metadata) was loaded into the Loupe Cell Browser, we then generated lists of marker genes for each of the identified clusters using the “Globally Distinguishing” feature. Marker genes were classified as those genes up-regulated in each cluster (compared to all other cells) with an adjusted p-value less than 0.1. Next we assigned tentative identities to each of these initial clusters by comparison of marker genes to available literature regarding markers of immune cells in teleost fish and other vertebrates. During this initial identification process, we identified multiple groups of cells with homology to the same major immune cell type (e.x. three clusters demonstrated patterns of expression indicative of neutrophils). Consequently, we condensed the initial 19 identified clusters into 8 major groups based on homology to known vertebrate immune cell types. We examined differential expression between sub-clusters within these major groups using the “Locally Distinguishing” feature in Loupe Cell Browser. Cluster identification and sub-cluster distinctions were confirmed by visual analysis of expression of major immune cell type markers in Loupe Cell Browser. Violin plots and heatmaps displaying patterns of expression across major group and sub-clusters within groups were generated in R using read count matrixes and cluster identity information (exported from Loupe Cell Browser). Relevant code can be found at: https://github.com/lfuess/scRNAseq.

### Comparative Analyses Across Populations

When comparing across populations, we assessed two hypotheses: **1)** relative abundance of immune cell types is variable across populations and **2)** expression patterns within each identified immune cell type are variable across populations. First, to identify differences in relative abundance of each of our 8 major immune cell types we performed independent, binomial general linear models for each cell type. Tukey’s post-hoc tests were used for pair-wise comparisons if significant differences were identified between populations (code can be found at: https://github.com/lfuess/scRNAseq). Second, to identify differences in gene expression patterns within each of our identified immune cell types, we again used the “Locally Distinguishing” feature in Loupe Cell Browser. Cells within each major group were subdivided by population, and then all possible pairwise comparisons of gene expression were conducted. Genes with adjusted p-values < 0.10 were identified as significantly differentially expressed. Relevant violin plots and heatmaps were generated in R using read count matrixes and cluster identity information (exported from Loupe Cell Browser). Relevant code can be found at: https://github.com/lfuess/scRNAseq.

### Sequence Alignment

In order to examine sequence divergence in the two identified copies of neutrophil marker gene, *nephrosin* (*npsn*), we conducted a multiple sequence alignment of both *npsn* transcripts from stickleback and the zebrafish *npsn* transcript sequence using the R package msa (72).

### Comparison to Past Analyses

We leveraged past transcriptomic analysis of stickleback head kidney to assess whether whole tissue-measured expression of putative markers identified here could be used as a reliable metric of relative cell type abundance. We specifically analyzed two past transcriptomic data sets: **1)** an analysis of laboratory-reared F1 fish from Roberts and Gosling Lake experimentally exposed to parasites (2), and **2)** an analysis of laboratory-reared F2 and backcrossed fish, the offspring of fish from experiment 1, experimentally exposed to parasites (59). For both of these datasets we had access to transcriptomic data detailing whole tissue expression of our putative cell markers, and flow cytometry data coarsely estimating granulocyte to lymphocyte relative abundance using forward and side-scatter gating. For each data set we examined correlation between normalized gene expression of putative markers and square root transformed frequency data for granulocytes or lymphocytes as appropriate.

Once we established that whole-tissue expression of putative cell markers was at least partially indicative of relative abundance of immune cell types, we then leveraged our newly identified cell markers to re-interpret three past transcriptomic studies of stickleback immunity: the two previously mentioned transcriptomic studies of F1 & F2/backcross fish to immune challenge (2, 59), and an additional study examining correlations between head kidney gene expression and gut microbiome composition (11). Specifically, we used chi squared tests to identify significant over-representation of markers of any given cell type within lists of genes significant differentially expressed as a result of traits of interest, or genes significantly correlated to microbial diversity/taxa of interest. Chi-squared tests were used to test for over-representation of each immune cell type within each list of genes independently.

## Acknowledgements

We gratefully acknowledge the contribution of Bill Flynn and Martine Seignon of the Single Cell Biology service as well as the Genome Technologies and Cyberinfrastructure high performance computing resources at The Jackson Laboratory for expert assistance with the work described in this publication. This work was supported by funding from NSF (IOS-1645170), the University of Connecticut (Startup to DIB), and an American Association of Immunologists Intersect Postdoctoral Fellowship to LEF.

## Competing Interests

The authors declare no competing interests.

## Notes

https://github.com/lfuess/scRNAseq

### Competing Interest Statement

The authors have declared no competing interest.

