## Supplementary Captions for "Single-cell RNA sequencing reveals micro-evolution of the stickleback immune system"

**Supplemental File Captions**

**Supplemental Table 1**: Description of the original 24 clusters, including top 5 marker genes for each cluster with significant markers, and condensed cell type assignment.

**Supplemental Table 2:** Results from GLM analysis and Tukey post-hoc tests for significant differences in cell type abundance between each of our three populations. Tukey post-hoc results are listed for each possible pairwise comparison with adjusted significance values.

**Supplemental Table 3:** Results of χ^2^ tests for over-representation of immune cell type markers generally, and each specific immune cell type, in lists of significantly differentially expressed genes from three previously published transcriptomic data sets. Proportion test results are also shown where χ^2^ test results are significant.

**Supplemental File 1:** Lists of significant marker genes for each of the original clusters. Significant marker genes are designated as those expressed at significantly higher levels in the cluster (padj < 0.1). Clusters with no significant marker genes omitted.

**Supplemental File 2:** Lists of significant marker genes for each of the eight condensed cell types. Significant marker genes are designated as those expressed at significantly higher levels in the cluster (padj < 0.1).

**Supplemental File 3:** Differential expression analysis results examining variation in gene expression of each cell type among three populations of fish. Significantly differentially expressed genes (padj < 0.1) for all pairwise comparisons are displayed.

**Supplementary Figure 1:** Loupe uMap demonstrating original cluster assignments for each cell. Each individual dot represented the expression profile of a single sequenced cell.

**Supplementary Figure 2:** Paired violin and Loupe uMap expression plots demonstrating patterns of expression for HC marker genes of interest. Plots display normalized (log-transformed) expression. Darker colors in cluster plots correspond to higher expression; each plot scaled independently.

**Supplementary Figure 3:** Paired violin and Loupe uMap expression plots demonstrating patterns of expression for neutrophil marker genes of interest. Plots display normalized (log-transformed) expression. Darker colors in cluster plots correspond to higher expression; each plot scaled independently.

**Supplementary Figure 4:** Paired violin and Loupe uMap expression plots demonstrating patterns of expression for APC marker genes of interest. Plots display normalized (log-transformed) expression. Darker colors in cluster plots correspond to higher expression; each plot scaled independently.

**Supplementary Figure 5:** Paired violin and Loupe uMap expression plots demonstrating patterns of expression for B-cells marker genes of interest. Plots display normalized (log-transformed) expression. Darker colors in cluster plots correspond to higher expression; each plot scaled independently.

**Supplementary Figure 6:** Paired violin and Loupe uMap expression plots demonstrating patterns of expression for RBC marker genes of interest. Plots display normalized (log-transformed) expression. Darker colors in cluster plots correspond to higher expression; each plot scaled independently.

**Supplementary Figure 7:** Paired violin and Loupe uMap expression plots demonstrating patterns of expression for platelet marker genes of interest. Plots display normalized (log-transformed) expression. Darker colors in cluster plots correspond to higher expression; each plot scaled independently.

**Supplementary Figure 8:** Paired violin and Loupe uMap expression plots demonstrating patterns of expression for fibroblast marker genes of interest. Plots display normalized (log-transformed) expression. Darker colors in cluster plots correspond to higher expression; each plot scaled independently.

**Supplementary Figure 9:** Paired violin and Loupe uMap expression plots demonstrating patterns of expression for NKC marker genes of interest. Plots display normalized (log-transformed) expression. Darker colors in cluster plots correspond to higher expression; each plot scaled independently.

**Supplementary Figure 10:** Heatmap displaying normalized (log transformed) expression of genes which were significantly differentially expressed between the two major types of B-cells. Heatmap and gene dendrogram generated using the R package, pheatmap.

**Supplementary Figure 11:** Alignment of predicted protein sequences of nephrosin from zebrafish, and the four predicted transcripts resulting from two nephrosin genes in the stickleback genome. Alignment generated using the R package msa.
